# Metabolic vulnerability is a target of the antineoplastic effect of breastfeeding

**DOI:** 10.64898/2026.03.03.709410

**Authors:** Edmund Charles Jenkins, Mrittika Chattopadhyay, Kristan Skriver Andersen, Soma Seal, Nicola Tavella, Joanne Stone, Sabine Heitzeneder, Crystal L. Mackall, Rachel Brody, Claus Oxvig, Doris Germain

## Abstract

Lactation is associated with a protective effect against breast and ovarian cancer as well as against cardiovascular diseases suggesting a systemic effect. Here, we show that the serum of lactating mice and breastfeeding mothers have targeted antineoplastic effects, while serum from virgin mice and matched post-partum but non-lactating women do not. The effect is specific to cancer cells expressing Pappalysin-A (PAPP-A), a target that is shared among diseases affected by breastfeeding. RNAseq revealed that lactating serum inhibits mitochondrial function and we found that PAPP-A alone lowers mitochondrial function, suggesting that lactation serum acts by exploiting the metabolic vulnerability of these cancer cells. Using serum proteomics, we identified corticotropin release factor (CRF) as being unique to serum of lactating women and we show that CRF alone mimics the mitochondrial and anti-tumorigenic effect of lactating serum. Blocking the CRF receptor, inhibits the protective effect of lactating serum. Since CRF has shown efficacy in the clinic in other settings, our findings raise the possibility to extend its use to mimic or enhance the protective effect of breastfeeding.

**Summary:** We show that the serum from lactating women has anti-cancer activity and used multi-omics approaches to identify corticotropin release factor as a peptide able to mimic the effect of lactating serum by targeting cells with low mitochondrial activity.

## INTRODUCTION

Breastfeeding is known to protect against breast cancer ^1,2^. Recent studies have implicated the role of increased T cells mediated immunity in the breast ^3,4^ as well as breast microbiota ^5^ as mechanisms to explain the local effect of breastfeeding in the breast. However, breastfeeding is also protective against ovarian cancers ^6–8^ as well as against atherosclerosis ^9–11^ and cardiovascular diseases ^12–14^. These observations raise the possibility that in addition to mechanisms acting locally in the breast, breastfeeding activates either additional mechanisms acting locally in other tissues or a systemic mechanism that acts on a target that is shared between these diseases.

Pregnancy-associated plasma protein A (PAPP-A) is a secreted protease that activate IGF signaling by mediating the degradation of IGFBP-4 and 5 ^15–20^. PAPP-A is frequently overexpressed in breast cancer ^21–26^ and ovarian cancer ^18,27–30^ as well as in atherosclerosis plaque ^31^ and is linked to cardiovascular diseases ^32–34^. PAPP-A is therefore a potential target that is shared among diseases that are targeted by the protective effect of breastfeeding.

In support of this hypothesis, we reported that while transgenic mice overexpressing PAPP-A in the mammary gland develop mammary tumors ^35^, lactation opposes the tumorigenic action of PAPP-A by inhibiting the effect of PAPP-A on post-partum involution of the mammary gland ^35^. The mechanism of how lactation protects against PAPP-A driven mammary tumor in these mice remained unknown.

In the current study, we focused on the potential systemic action of lactation against breast cancer. We collected serum from lactating mice as well as breastfeeding mothers and tested their effect on the growth of cancer cells. Our data demonstrate an anti-tumorigenic effect of serum from lactating mothers and that this effect is specific toward cancer cells expressing PAPP-A. We further identified corticotropin release factor (CRF) as the mediator of this effect.

## RESULTS

### Serum from breastfeeding women suppresses the growth of PAPP-A+ cancer cells

To test the effect of lactating serum, since we previously shown that lactation opposes the oncogenic action of PAPPA in mice^35^, we selected 3 breast cancer cell lines that show either no detectable level of PAPP-A (MCF7), elevated level of PAPP-A due to overexpression (MCF7-PAPP-A) or endogenous level of PAPP-A (Hcc70) (Fig. 1A). Blood was collected from 3 virgin female mice (non-lactating) and 4 lactating female mice, serum was isolated and added to culture media at a final concentration of 10% in absence of fetal calf serum. The resulting media was added to 3D cultures of either MCF7, MCF7-PAPP-A or Hcc70 breast cancer cells and colonies size determined. We found that the size of the colonies of MCF-7 cells was the same when treated with serum derived from non-lactating (NL) or lactating (L) females (Fig. 1B, C). However, the colonies of MCF7-PAPP-A (Fig.1D, E) and Hcc70 (Fig. 1F, G) cells were significantly smaller upon incubation with the serum from lactating females compared to serum from non-lactating virgin mice. This result suggests an anti-proliferative effect of lactating serum that is specific to cells expressing PAPP-A. To determine if this effect of lactating serum is also observed in humans, we collected blood and isolated serum from 30 women at their 6 weeks post-partum visit. Of these 30 women, 9 did not lactate and 21 lactated and were still lactating at their 6 weeks visit (Fig. 1H). Additionally, since pregnancy involves multiple hormonal and other physiological changes independently of lactation, we also obtained blood and isolated serum from 10 pre-menopausal women who never had children (Fig. 1H, nulliparous). The effect of each serum was tested on 3D culture of MCF7 or MCF7-PAPP-A cells and we found that, as observed in mice (Fig. 1B-E), no effet was observed on MCF7 cells (Fig. 1I) but the growth inhibitory effect of lactating serum was on MCF7-PAPP-A cells (Fig. 1J). To verify that the effect is specific to lactating serum, we also tested the effect of serum from nulliparous women on MCF7-PAPP-A cells and found no effect (Fig. 1J). To further test the correlation between expression of PAPP-A and the effect of lactating serum, we tested 3 additional cells lines that do not overexpress PAPP-A and found that lactating serum has no effect on these cells (Suppl. Fig. 1).

**Figure 1:**
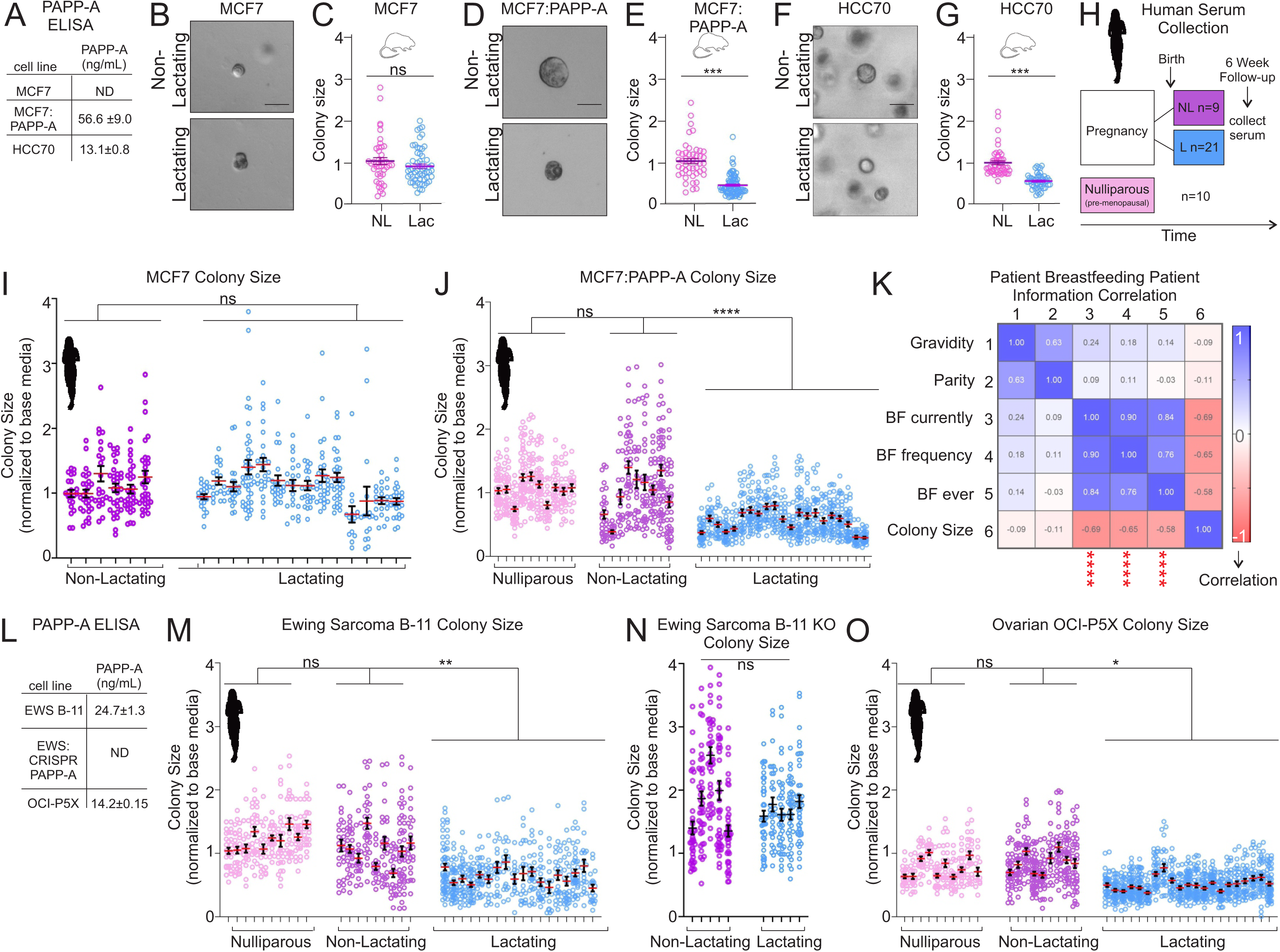
Serum from lactating women suppresses the growth of cancer cells expressing PAPP-A. A) ELISA of PAPP-A levels detected in media of indicated cell lines. ND= not detected. B) Representative image of 3D cell culture of MCF7 cells grown in media supplemented with 10% serum from non-lactating or lactating mice. C) Quantification of B. D) Representative image of 3D cell culture of MCF7-PAPP-A cells grown in media supplemented with 10% serum from non-lactating or lactating mice. E) quantification of D. F) Representative image of 3D cell culture of HCC70 cells grown in media supplemented with 10% serum from non-lactating or lactating mice. G) Quantification of F. H) Schematic representation of cohorts of women who donated blood. NL= non-lactating. L= Lactating. NP= Nulliparous. I) Quantification of 3D colony size of MCF7 cells grown with serum from non-lactating or lactating women. J) Quantification of 3D colony size of MCF7-PAPP-A cells grown with serum from nulliparous, non-lactating or lactating women. K) Person Correlation analysis of patient clinical information of MCF7-PAPP-A colony size. L) ELISA of PAPP-A levels detected in media of indicated cell lines. M) Quantification of 3D colony size of Ewing Sarcoma cells grown with serum from nulliparous, non-lactating or lactating women. N) Quantification of 3D colony size of Ewing Sarcoma PAPP-A Knock-out cells grown with serum from non-lactating or lactating women. O) Quantification of colony size of Ovarian OCI-P5X 3D cells grown with serum from nulliparous, non-lactating or lactating women. Statics: Student’s (panels C, E, and G) or nested (panels I, J, M, N, O) t-test. *p<0.05, **p<0.005, ***p<0.0005, ****p<0.00005. Scale bars = 50um

To determine if correlation exists between the history of breastfeeding and the anti-cancer activity of the lactating serum, we performed a multi-variate analysis and found a strong negative correlation between colony size and breastfeeding, independently of the frequency (Fig.1K).

To further clarify the apparent specificity of the effect on cancer cells expressing PAPP-A, we expanded our analysis to Ewing Sarcoma (EWS B-11) and ovarian cancer (OCI-P5X) cancer cells, which have been reported to endogenously overexpress PAPP-A ^18,21,27–30,36–38^ and included a clone of EWS B-11 where PAPP-A gene has been deleted by CRISPR (EWS CRISPR PAPPA). First, we established PAPP-A levels by ELISA and found that they express higher or similar level as the Hcc70 breast cancer cell line (Fig. 1L). We repeated the 3D colonies assay using EWS B11 cells and found that the lactating serum reduced their growth relative to serum from non-lactating women (Fig. 1M) and confirmed that serum from nulliparous women had no effect. To confirm the specificity of the effect against cells expressing PAPP-A, we tested non-lactating and lactating serum on EWS CRISPR PAPPA cells and found that in absence of PAPP-A the inhibitory effect of lactating serum is no longer observed (Fig. 1N). Inhibition of colony formation by lactating serum was also confirmed using endogenously overexpressing PAPP-A ovarian cancer cells OCI-p5X (Fig. 1O).

These results suggest that the protective effect of lactation against cancer cells is mediated systemically through factors present in the serum of breastfeeding women and that the effect is specific to cancer cells expressing PAPP-A.

### RNAseq analysis identifies mitochondrial function as the target of serum from breastfeeding women

To interrogate the mechanism by which lactation serum affects the growth of PAPP-A driven cancer cells specifically, we performed RNA seq on MCF7-PAPP-A 3D colonies exposed either to serum derived from lactating and non-lactating post-partum women. Principal component analysis revealed a strong clustering of samples from the lactating and non-lactating groups derived from the 3D colonies (Fig. 2A). We then performed a search for pathway enrichment terms from RNAseq derived from the 3D cultures and found that mitochondrial related pathways and cell cycle are down-regulated by lactating serum, while cell death and DNA damage pathways are increased (Fig. 2B). To better define these terms, we performed pathway analysis and found that the mitochondria related pathways are mainly linked to the down-regulation of the electron transport chain and ATP generation, while the up-regulated pathways are associated with oxidative stress-induced senescence, cell death and DNA damage (Fig. 2C). Since mitochondrial pathways were the most frequent and significant pathways altered by lactating serum, the genes leading to these pathways were identified (Fig. 2D) and heat map analysis confirmed that they are significantly down-regulated in organoids exposed to lactating serum compared to non-lactating serum (Fig. 2E). To gain further insight into how widely mitochondria-related genes are affected by lactating serum, we also compiled genes from the MitoCarta (Suppl. Fig. 2) and found that while they are not as strongly downregulated as those identified by RNAseq, this analysis indicates a global down regulation of mitochondria-related genes (Fig. 2F).

**Figure 2:**
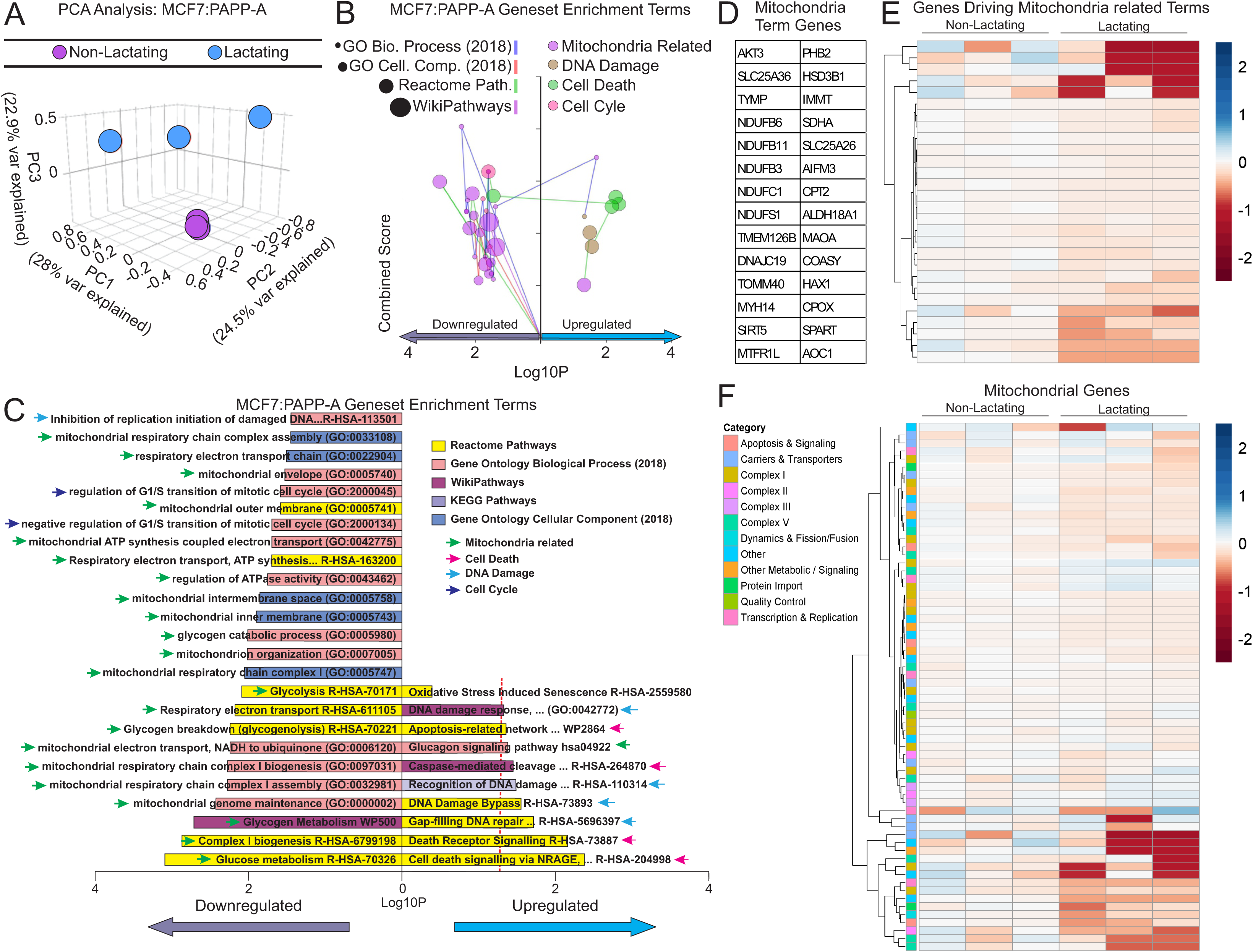
RNAseq analysis identifies suppression of mitochondrial activity as a main pathway that is altered by serum from lactating women. A) Principle component analysis of RNAseq data from 3D colony of MCF7-PAPP-A cells incubated with 10% serum from non-lactating of lactating women (n=3 per group). B) Summary of GSEA of the top 500 differentially regulated genes comparing the non-lactating and lactating groups using the ENRICHR platform. C) Most significant term hits across indicated databases of the top 500 differentially expressed genes between the non-lactating and lactating groups. D) List of genes driving the term hits in panel C. E) Heatmap of genes in panel D between 3D culture exposed to non-lactating or lactating serum. F) Heatmap of gene expression of genes in panel E and additional mitochondrial genes selected from MitoCart3.0.

Next, to determine if the effect of lactating serum is also observed *in vivo,* we performed xenografts using MCF7-PAPP-A cells. For this experiment, due to the limitation in the amount of material available, we pulled the serum from 5 breastfeeding women and 5 non-lactating women and once tumors were detected in all mice, we injected serum intraperitoneally once a day for one week and measured tumor volumes over time. We found at the endpoint of the experiment that MCF7-PAPP-A tumors were significantly smaller in size and weight in the lactating group (Fig. 3A, B, C) and their growth rate significantly slower over time (Fig. 3D). We then performed RNAseq on the tumors exposed to either non-lactating or lactating serum. As observed with the organoids, we found a strong clustering of xenografts injected with lactating and non-lactating serum using principal component analysis (Fig. 3E) and that pathway enrichment terms derived from RNAseq are associated with mitochondria pathways and cell cycle and are down-regulated by lactating serum, while cell death and DNA damage pathways are increased (Fig. 3F). The number of genes associated with mitochondria pathways was even larger in xenografts compared to 3D culture exposed to lactation serum (Suppl. Fig. 3) and heat map analysis also showed a strong clustering and downregulation of these genes in lactating serum treated tumors *in vivo* (Fig. 3G). Broader analysis using all genes of the MitoCarta also demonstrated down-regulation of mitochondrial related genes (Fig. 3H). Finally, we compared the pathways identified in 3D culture and xenografts and found a significant overlap in pathways related to mitochondria (Fig. 3I) despite the fact the multiple differences exist between *ex vivo* and *in vivo* experimental conditions.

**Figure 3:**
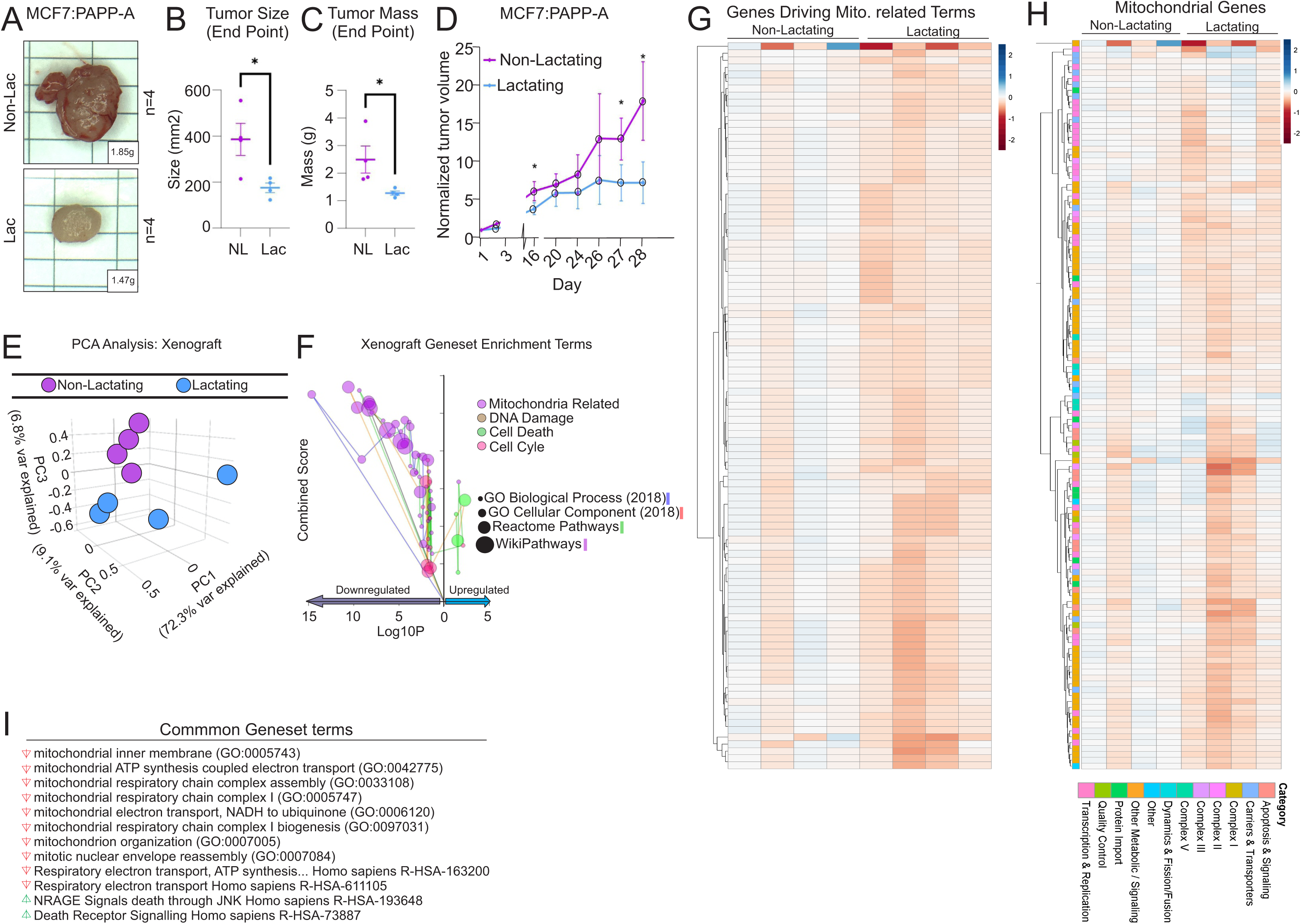
Serum from lactating woman suppresses the growth of MCF7-PAPP-A xenografts and expression of mitochondrial genes. A) Representative images of MCF7-PAPP-A xenografts treated with non-lactating (NL) or lactating (L) serum. Grid 0.5×1cm. B) Graph of size of MCF7-PAPP-A xenografts exposed to NL and L serum. C) Graph of weight of MCF7-PAPP-A xenografts exposed to NL or L serum. D) Growth curve of MCF7-PAPP-A xenografts exposed to NL and L serum. E) Principal component analysis of RNAseq from MCF7-PAPP-A xenografts from mice treated with NL and L serum. F) Summary of GSEA of the top 500 differentially regulated genes comparing the non-lactating and lactating groups using the ENRICHR platform. G) Heatmap of genes driving the term hits in panel F. H) Heatmap of gene expression of genes in panel G and additional mitochondrial genes selected from MitoCart3.0. I) Geneset terms that were significantly enriched in both 3D cell culture and xenograft experiment.

Collectively, these analyses indicate that mitochondria related pathways are a central target of lactating serum.

### Serum from breastfeeding women suppresses mitochondrial function leading to DNA damage and Senescence

Since the RNAseq from both *ex vivo* and *in vivo* analyses point to a strong effect of the lactating serum on ATP production by the mitochondria, we next aimed at measuring mitochondrial activity using Seahorse upon exposure to non-lactating and lactating serum in both MCF7 and MCF7-PAPP-A breast cancer cells (Fig. 4A). We found that basal respiration (Fig. 4B), respiration coupled ATP production (Fig. 4C) and maximal respiration (Fig. 4D) were all significantly lower in MCF7-PAPP-A cells treated with serum from breastfeeding women relative to those treated with serum from non-lactating post-partum women. However, no difference in mitochondrial activity was detected by Seahorse in MCF7cells exposed to lactating versus non-lactating serum (Fig. 4A-D).

**Figure 4:**
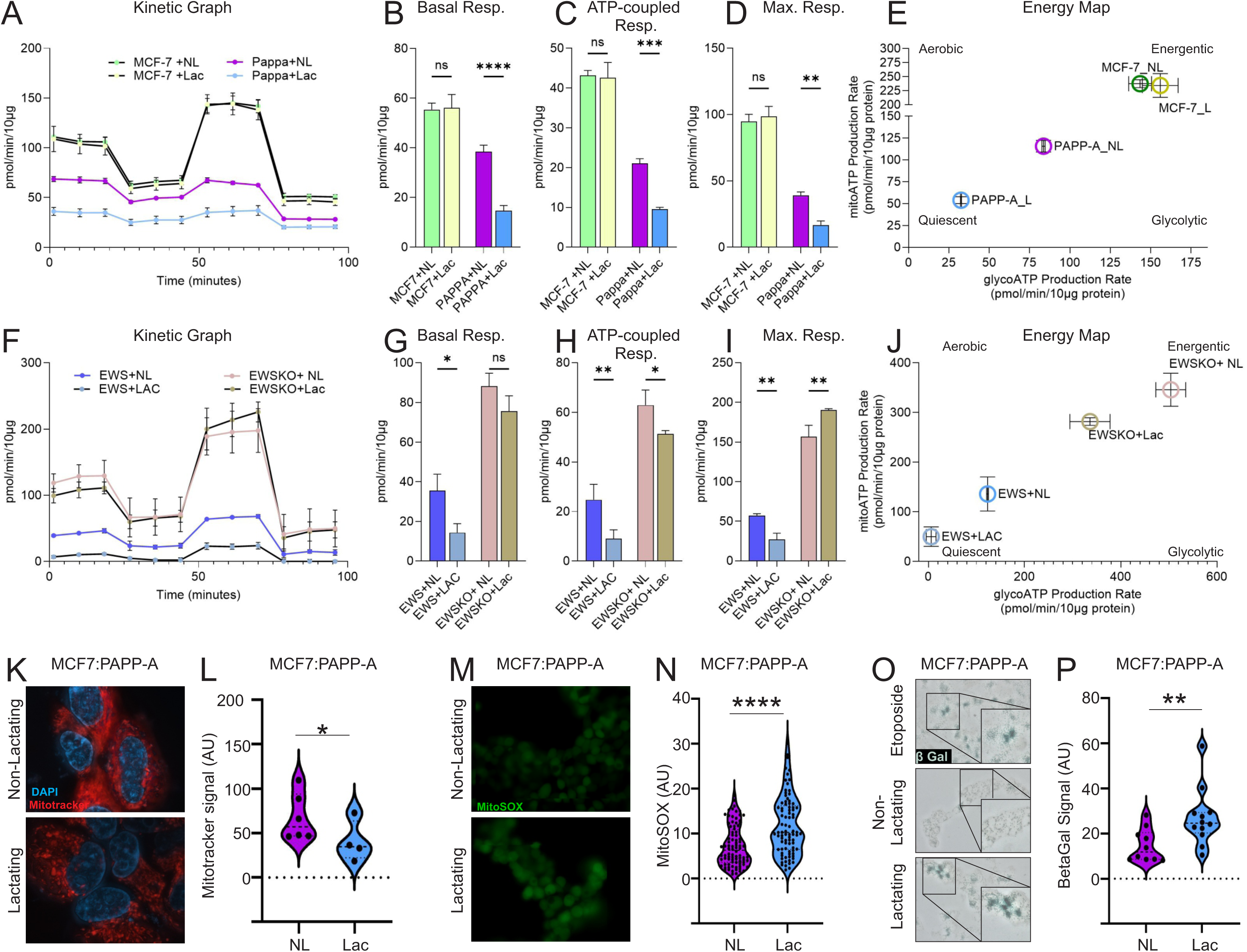
Serum from lactating women suppresses mitochondrial function in PAPP-A+ cells. A) Kinetic graph of MCF7 and MCF7-PAPP-A cells grown with 10% serum from non-lactating (NL) or lactating (Lac) women. B) Basal Respiration, C) ATP-coupled Respiration, and D) maximal respiration of MCF7 and MCF7-PAPP-A cells grown with 10% serum from NL or Lac women. E) Energy map of MCF7 and MCF7-PAPP-A (PAPP-A) cells grown with 10% serum from NL or Lac women. F) Kinetic graph of Ewing Sarcoma (EWS) and Ewing Sarcoma PAPP-A knock-out cells (EWSKO) grown with 10% serum from NL or Lac women. G) Basal Respiration, H) ATP-coupled Respiration, and I) maximal respiration of Ewing Sarcoma (EWS) and Ewing Sarcoma PAPP-A knock-out cells (EWSKO) grown with 10% serum from NL or Lac women. J) Energy map Ewing Sarcoma (EWS) and Ewing Sarcoma PAPP-A knock-out cells (EWSKO) grown with 10% serum from NL or Lac women. K) Representative images of MCF7-PAPP-A cells treated with serum from NL or Lac women and stained with Mitotracker. L) Quantification of K. M) Representative images of MCF7-PAPP-A cells treated with serum from NL or Lac women and stained with MitoSOX. N) Quantification of M. O) Representative images of MCF7-PAPP-A cells treated with Etoposide (positive control), or serum from NL or Lac women and stained with X-gal to assess Beta Galactosidase activity. P) Quantification of O. Significance was determined using student’s t-test. *p<0.05, **p<0.005, ***p<0.0005, *****p<0.00005.

Energy map, which evaluate both ATP production generated by glycolysis and by the mitochondria, confirmed that MCF7 cells have elevated mitochondrial activity independently of being treated with lactating or non-lactating serum (Fig. 4E). In contrast, MCF7-PAPP-A cells exposed to lactating serum had the lowest mitochondrial activity suggesting senescence (Fig. 4E). Interestingly, these results also revealed that overexpression of PAPP-A alone correlated with lower mitochondrial activity since the activity in MCF7-PAPP-A cells exposed to non-lactating serum is lower than MCF7 cells exposed to non-lactating serum (Fig. 4A-E). To further, test this observation, we used the Ewing sarcoma cell line EWS B11, where PAPP-A is overexpressed endogenously and its CRISPR counterpart, where PAPP-A has been deleted. First, as observed in the MCF7-PAPPA model, we found that the mitochondrial activity of EWS B11 cells is decreased by lactating serum (Fig. 4F, G, H, I) but not in EWS B11 cells where PAPP-A is deleted (Fig. 4F, G, H, I). Additionally, this analysis confirmed that mitochondrial function is lower when PAPP-A is expressed (Fig. 4F, blue line) compared to PAPP-A KD cells (Fig. 4F, pink line). The effect of PAPP-A on the activity of the mitochondria is in agreement with 1) the finding that mitochondrial function is increased in the PAPP-A knockout mice ^39^, 2) the link between PAPP-A and the interaction between the endoplasmic reticulum and mitochondrial function ^40^ and 3) several studies demonstrating a link between IGF signaling and mitochondrial function ^41–45^. Therefore, our finding adds to this growing number of reports.

Energy map analysis indicated the same pattern as observed in the MCF7 breast cancer model, where PAPP-A expression confers lower mitochondrial function and this is further exacerbated upon exposure to lactating serum (Fig. 4J).

To obtain independent assessment of mitochondria function, we used Mitotracker red staining and found a decrease in staining indicating lower mitochondrial inner membrane potential (Fig. 4K, L). Further, since the RNAseq data indicated an increase in oxidative stress and DNA damage pathways, we also measured the level of mitochondrial oxygen reactive species (ROS) using MitoSOX staining and found that the level of ROS is increased in MCF7-PAPP-A cells treated with lactating serum relative to non-lactating serum (Fig. 4M, N). To assess DNA damage, we used staining with phosphorylated gamma histone H2A (p-γH2A), which marks sites of DNA damage and found a significant increase in p-γH2A staining (Suppl. Fig. 4). Since the mitochondria is the hub of several cell death pathways ^46^, we next monitored the potential induction of apoptosis. However, since we found no significant difference in the cleavage of caspase 3, we focused on senescence using staining for β-gal and found a significant increase in β-gal staining (Fig. 4O, P).

These results validate the data obtained by RNAseq and indicate that serum from breastfeeding women inhibits the growth of PAPP-A+ cancer cells by exploiting their mitochondrial vulnerability leading to mitochondrial shutdown, increased ROS and senescence.

### Corticotropin-release factor mediates the antineoplastic effect against PAPP-A+ breast cancer cells

To identify the factor responsible for the effect of serum from breastfeeding women, we performed untargeted proteomics on all serum of lactating (n=21) and non-lactating (n=9) age-matched post-partum women. The samples were screened for the presence a total of approximately 11, 000 proteins (Fig. 5A). This analysis revealed 359 proteins were significantly differentially regulated between groups (Fig. 5B). Of these, 209 were higher in the serum from lactating women. Using the Human Protein Altas, we determined that 16 of those proteins are known to be secreted into serum (Fig. 5B, C). Among those 16 proteins, 2 (prolactin and corticotropin-release factor (CRF) were of particular interest given their association in the literature with the terms “lactation”, “mitochondria”, and “breast cancer” (Fig. 5D), Since prolactin is specific to lactation, we view its detection by proteomics as an internal control validating our dataset. Additionally, prolactin is reported to promote the transcription of CRF ^47^ and is functionally linked to its action ^48–50^. CRF, while produced by the placenta during pregnancy ^51^, is regulated through the hypothalamus-pituitary-adrenal (HPA) axis during lactation ^52,53^, is known to disrupt mitochondrial morphology and function ^54–57^ and additionally has reported anti-cancer effects ^58–63^, although a link between the effect of CRF on mitochondria and its anti-cancer effect has never been described. For these reasons, and the fact that CRF was significantly up-regulated (Suppl. Fig. 5), we focused on CRF. Therefore, to further validate that the detection of CRF is specific to lactating serum, we used CRF ELISA to monitor the level of CRF in the serum of nulliparous women using lactating serum as positive control. This analysis showed that CRF is not detected in the serum of nulliparous women (Fig. 5E), while it is detected in lactating serum, in agreement with the proteomics data.

**Figure 5:**
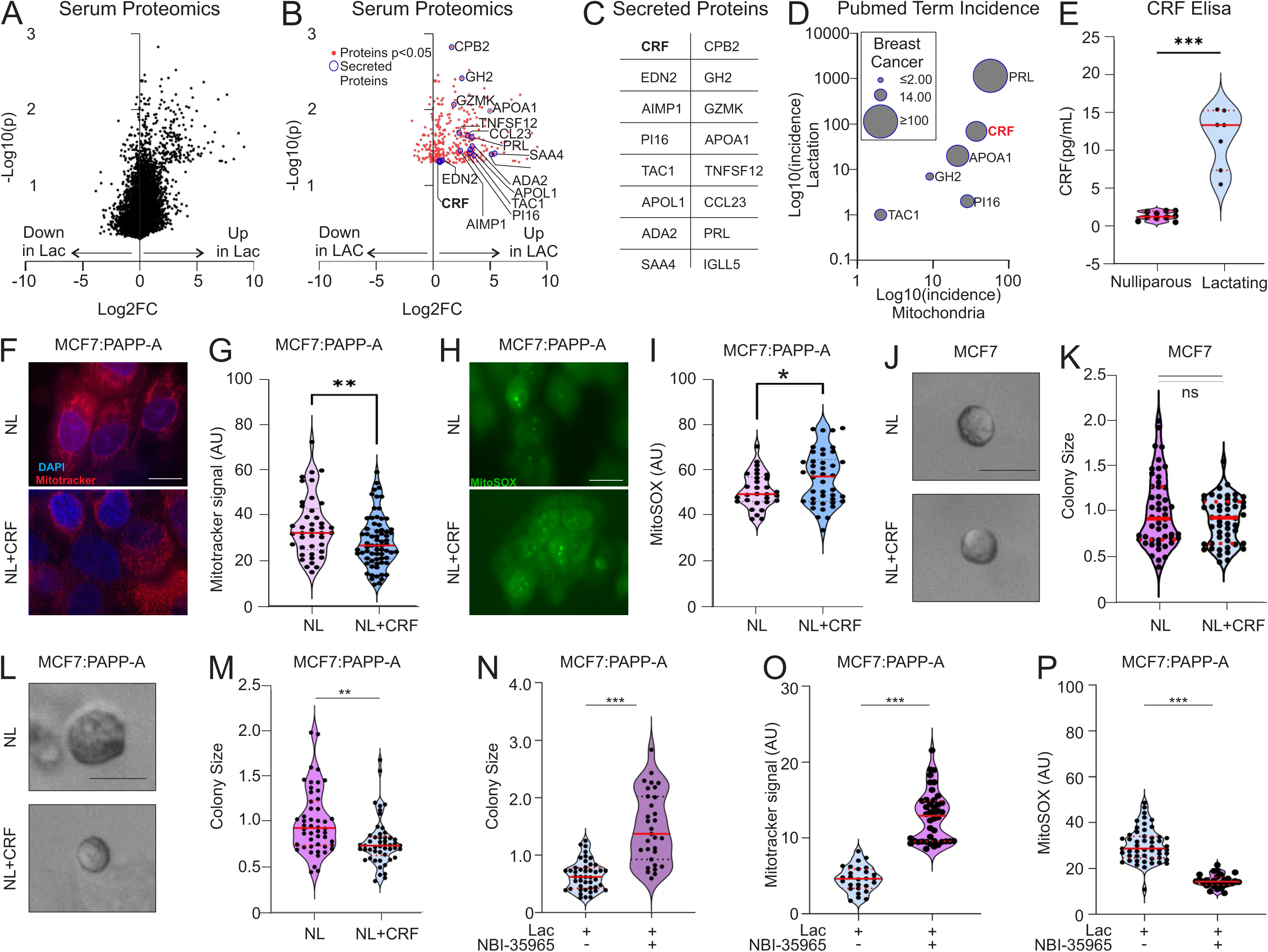
Serum proteomics identifies CRF as the driver of the effect of serum from lactating women. A) Volcano plot of proteomic data comparing serum from non-lactating (n=9) and lactating women (n=21). B) Graph of serum proteins significantly up-regulated in lactating serum (n=21) compared to non-lactating serum (n=9). Secreted proteins are indicated with blue circles. C) Table of secreted proteins from panel B. D) Graph of ranking of secreted proteins based on term incidence using Pubmed search combining “Mitochondrial”, “Lactation”, and “Breast Cancer” as search terms. E) ELISA for CRF in serum from nulliparous (n=10) or lactating (n=7) women. F) Mitotracker staining of MCF7-PAPP-A cells cultured with serum from non-lactating women plus or minus 50nM CRF. Scale bars = 25um G) Quantification of F. H) MitoSOX staining of MCF7-PAPP-A cells cultured with serum from non-lactating women plus or minus 50nM CRF. Scale bars = 25um. I) Quantification of H. J) 3D cell culture of MCF7 cells cultured with serum from non-lactating women plus or minus 50nM CRF. Scale bars = 50um. K) Quantification of J. L) 3D cell culture of MCF7-PAPP-A cells cultured with serum from non-lactating women plus or minus 50nM CRF. M) Quantification of L. Scale bars = 50um. N) Quantification of 3D cell culture of MCF7-PAPP-A cells cultured with serum from lactating women plus or minus the CRF inhibitor, NBI-35965 (50nM). O) Quantification of Mitracker or MitoSOX (P) staining for MCF7-PAPP-A cells cultured with serum from lactating women with or without the CRF receptor inhibitor NBI-35965 (50nM). Statistical significance was determined by student’s t-test. *p<0.05, **p<0.005, ***p<0.0005.

CRF is produced as a result of proteolytic cleavage from a precursor protein through the secretory pathway and released as an active hormone of 41 amino acids. Corticorelin acetate is the synthetic form of CRF and has been used clinically in the setting of brain cancer ^64^. To determine if CRF alone can mimic the effect of lactating serum on breast cancer cells, we repeated the analyses using Mitotracker and MitoSOX staining and colony formation in 3D culture. We found that adding CRF to non-lactating serum reduces Mitotracker staining (Fig. 5F, G) and increases staining with MitoSOX (Fig. 5H, I) indicating lower mitochondrial inner membrane potential and increased oxidative stress respectively. CRF has no effect on colony formation of MCF-7 cells (Fig. 5J, K) but significantly reduced the growth of MCF7-PAPP-A breast cancer cells (Fig. 5L, M). To further validate the effect of CRF, we reasoned that since the action of CRF requires the binding to the CRF receptor, blocking binding to its receptor should abolish its effect. We therefore compared the effect of lactating serum with and without the CRFR inhibitor, NBI-3565. We found that inhibiting the action of CRF abolishes the effect of lactating serum on MCF7-PAPP-A cells colony formation (Fig. 5N), Mitotracker (Fig. 5O) and MitoSOX (Fig. 5P).

We next addressed the effect of CRF on mitochondrial function by Seahorse using lactating and non-lactating serum, as an internal control. We found that CRF reduces mitochondrial function at the same level of lactating serum (Fig. 6A, B, C, D), while it had no effect on MCF7 cells (Fig. 6A,B, C, D). Analysis by energy map showed that the effect of treatment with lactating serum or CRF overlap (Fig. 6E) and both push cells toward senescence, while CRF treatment in MCF7 cells had no impact (Fig. 6E). Conversely, adding the CRF receptor inhibitor NBI-3965 to lactating serum, inhibited its effect on the activity of the mitochondria in MCF7-PAPP-A cells (Fig. 6F, G, H, I, J). Finally, to complement these results, we tested the effect of CRF on the Hcc70 breast cancer cells, which overexpressed PAPP-A at endogenous level (Fig. 1A), using Seahorse and colony formation. We found that CRF also reduces mitochondrial function in these cancer cells at the same level as lactating serum (Fig. 6K, L, M, N, O), an effect that was also mirror by energy map analysis (Fig. 6P). Further, addition of CRF to non-lactating serum reduced colony formation in these cells (Fig. 6P).

**Figure 6:**
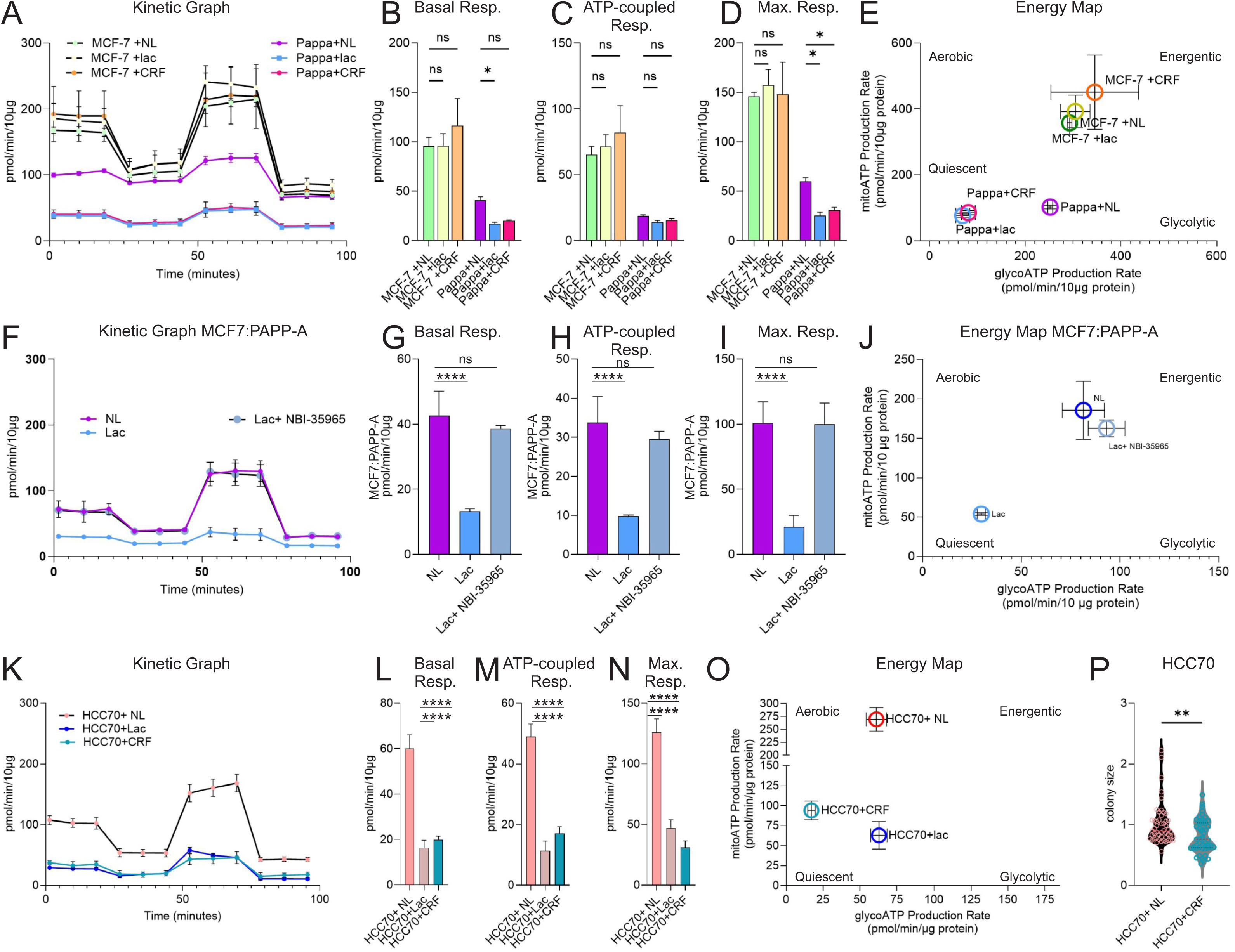
CRF suppresses mitochondrial function in PAPP-A+ cells. A) Seahorse analysis of MCF7 and MCF7-PAPP-A (PAPP-A) cells grown with 10% serum from non-lactating (NL) plus or minus CRF or lactating (Lac) women. Graphs of B) Basal respiration, C) ATP-coupled Respiration, D) maximal respiration and E) Energy Map (Basal levels) generated from analysis in A. Significance was determined by student’s t-test. F) Seahorse analysis of MCF7-PAPP-A (PAPP-A) cells growth with 10% serum from non-NL (internal control) or lactating (Lac) women plus or minus CRF receptor inhibitor NBI-35965 (50nM). Graphs of G) Basal respiration, H) ATP-coupled Respiration, I) Maximal respiration and J) Energy map, generated from analysis in F. K) Seahorse analysis of HCC70 cells grown with 10% serum from either non-lactating (NL) plus or minus CRF or lactating (Lac) women. Graphs of L) Basal respiration, M) ATP-production coupled Respiration, N) Maximal respiration and O) Energy map generated from analysis in K. P) Graph of colony size of Hcc70 breast cancer cells incubated in non-lactating serum with and without CRF. Significance was determined by student’s t-test. Statistical significance was determined by student’s t-test. *p<0.05, **p<0.005, ***p<0.0005.

These results support the conclusion that CRF in the serum of breastfeeding women plays a significant role in its antineoplastic effect. Our data however suggest that the effect of CRF is mainly restricted to PAPP-A expressing cancer cells due to their mitochondrial vulnerability.

## DISCUSSION

Pregnancy-associated plasma protein A (PAPP-A) was isolated in the blood of pregnant women in 1970 ^65^. Nearly 20 years later, it was identified as a protease with specificity toward the insulin-like growth factor binding proteins 4 and 5 (IGFBP-4, 5) ^15–20^. During pregnancy, PAPP-A is produced by and bound to the surface of placental trophoblasts but its role in the placenta remains unclear. However, considering that PAPP-A promotes proliferation ^17^ and immune evasion in cancer ^36^, it is tempting to speculate that the roles of PAPP-A in the placenta are similar as those it plays in cancer, since several parallels exist between placenta and cancer in terms of proliferation and invasion of tissue ^66–70^.

Our study shows that in both mice and human, when applied directly on established cancer cells, the serum of lactating females selectively inhibits the growth of PAPP-A positive cancer cells. PAPP-A is overexpressed across breast cancer sub-types including TNBC ^25^. Likewise, while breastfeeding is protective against all sub-types, some studies suggest a more potent effect against triple negative breast cancer ^71,72^. However, given that this sub-type is more frequent in young women than in post-menopausal women more studies are needed to confirm an association, if any, between breastfeeding and breast cancer sub-types. The results on the systemic effect of lactation presented in the current study highlight that a direct association between specific breast cancer sub-types and breastfeeding may be more complex than originally thought and that both local ^3,4^ and systemic effects may impact such association.

RNAseq of gene expression of breast cancer cells treated with serum from lactating women identified mitochondrial function as a central pathway targeted by lactating serum. Collectively, since we discovered that expression of PAPP-A leads to lower mitochondrial activity, these findings suggest that the specificity of the effect of lactating serum on cancer cells expressing PAPP-A acts by exploiting their metabolic vulnerability. This link between mitochondrial dysfunction and the antineoplastic effect of lactation serum is further supported by the identification of CRF by serum proteomics, since CRF is known to disrupt mitochondrial morphology and function ^54–57^ and additionally has reported anti-cancer effects ^58–63^. Therefore, our data unify a body of literature on independent observations and offer a functional link between the protective effect of breastfeeding against cancer and the effect of CRF on mitochondrial function.

Considering that PAPP-A is expressed at the surface of placental trophoblasts during pregnancy, our findings raise the possibility that under normal physiological conditions, the production of CRF through the HPA axis after pregnancy may ensure that any residual trophoblasts expressing PAPP-A during pregnancy are eliminated after pregnancy. If so, lactation may be a natural protection against gestational trophoblastic neoplasia ^73,74^. In addition to this physiological function however, our current data suggest that if any PAPP-A+ breast or ovarian cancer cells are present, lactation would inadvertently also allow their elimination and contribute to the protective effect of lactation against these cancers.

Further, since inhibition of PAPP-A eliminates atherosclerosis plaques ^31,32,75,76^, our findings may also explain the protective effect of breastfeeding against atherosclerosis ^9–11^

As the synthetic form of CRF (cortecorelin acetate) has already been used in the clinic ^64,77,78^, our study raises the possibility to expand its use not only to PAPP-A+ breast and ovarian cancers but potentially also to other clinical settings including as a protective agent against cardiovascular diseases and for the treatment of Ewing Sarcoma, a rare type of cancer affecting mainly affecting adolescents and young adults.

Caution about the use of CRF must however be acknowledged, since production of CRH in combination with other neuropeptides during chronic stress can contribute to cardiovascular diseases ^79,80^. Therefore, dosing and time of exposure must be taken into consideration to maximize the beneficial effect described here, while limiting the potential harmful effects associated with chronic stress.

In conclusion, our study suggests that in addition to the local effect of breastfeeding in the breast, the protective effect of lactation also acts through a systemic CRF-mediated mechanism that drives the elimination of cells that express PAPP-A, a target that is shared among conditions known to benefit from the protective effect of breastfeeding.

## METHODS

### Cell lines and 3D culture

MCF7, MCF7 cell overexpressing PAPP-A (MCF7-PAPP-A) ^81^, EWS B11 and EWS CRISPR-PAPP-A cells were grown and maintained in DMEM F-12 supplemented with 10% fetal calf serum and penicillin/streptomycin. OCI-P5X were cultured and maintained in 3D culture was performed similarly to as previously described ^82^. Briefly, 500-750 cells were cultured in 50µL of growth factor reduced Matrigel (BD sciences) and 150 µL growth media. Cultures were maintained in growth media containing 10%FCS for 24 hours before being washed three times in serum free media and then cultured in growth media supplemented with 10% serum from lactating or virgin mice, or lactating or non-lactating women. Cultures were allowed to grow for an additional 48hrs before being imaged.

### Image quantification

3D cultures were imaged using an Axio Observer Live imaging system (Zeiss) under identical imaging conditions. Colony size was determined by measuring two-dimension area of each 3D spheroid and weas performed in blinded manner, manually using ImageJ. Colony size was normalized to the mean colony size in the media containing 10% FCS. For each condition, 30-50 individual colonies were measured per individual serum (human n=30 or 40 when nulliparous also used, mice = 7).

### Statistics

Statistical significance was defined as a p value below 0.05 *p < 0.05, **p < 0.01, ***p < 0.001, ****p < 0.0001. Data displayed as mean ± the standard error of the mean (SEM). Statistical analyses were performed using GraphPad Prism Software 10. Statistical significance was determined using Student’s t-test with Welch’s correction (for two conditions) or one way ANNOVA followed by a Bonferroni post analysis. For proteomics analysis, the log2FC for each protein was calculated for each patient. Student’s t-test was performed between groups was performed to determine proteins that was differentially regulated.

### Human Blood collection

Human blood was collected under a protocol approved by the IRB at Mount Sinai. Blood was collected from human volunteers 6 weeks postpartum. Women were considered to be lactating if they were still lactating at their 6 weeks post-partum visit and considered non-lactating if they never lactate (7) or lactated for three days or less (2).

### Xenograft establishment

All animal experiments were performed under an IACUC approved protocol at Mount Sinai. Xenografts were established as previously described ^81,82^. 3-4 month old nude mice (Jackson Laboratories) were injected with 1 million MCF7-PAPP-A in 50:50 growth factor reduced Matrigel and growth media in the fourth inguinal mammary glands. A 60-day release 0.5 mg β-estradiol 17-acetate pellet (Innovative Research of America, #SE-271) was injected one week prior to xenograft establishment. Xenografts were allowed to establish for two weeks before treatment with serum from either lactating of non-lactating women. 50uL of patient serum was delivered by IP injection daily for 7 days.

### PAPP-A Elisa

PAPP-A levels were measured similarly to as previously described ^81^. 5 million cells were seeded and allowed to adhere for 24 hours in growth media. Media was then replaced with serum free media and allowed to be conditioned for an additional 24 hours. The Quantikine PAPP-A ELISA kit (R&D Systems, # DPPA00) was then used following the manufacturer’s protocol to determine the concentration of PAPP-A in conditioned media.

### RNA sequencing

Freshly isolated and snap frozen xenograft tissue (n=4) or snap frozen 3D cultures (N=3) were collected and sent to Azenta (Genewiz, South Plainfield, NJ) where they were processed for bulk paired end RNA sequencing. Raw FASTQ files were then aligned and normalized using either Biojupies ^83^ or BioStars on the Galaxy platform. Geneset enrichment analysis was performed across multiple databases using the Biojupies platform. Primary FASTQ files have been deposited and GEO numbers available upon publication.

### Seahorse

A Seahorse XF24 Extracellular Flux Analyzer (Agilent/Seahorse Biosciences, North Billerica, MA) was used to assess mitochondrial respiratory function. 25,000–50,000 cells per well were seeded, depending on cell size, to ensure formation of a uniform monolayer. Cells were plated on Seahorse XF24 plates in DMEM/F12 medium supplemented with 10% FBS for wells designated for CRF treatment, or serum-free DMEM/F12 for wells designated for treatment with human or mouse serum. Four to five hours after plating, cells were treated with 10% non-lactating (NL) serum, 10% lactating (L) serum, or NL serum plus 50nM CRF and incubated for 24 hours. For inhibition of the CRF receptor, cells were incubated with the CRF1 inhibitor (NBI-35965; Cayman Chemical Cat #18524), 50nM one hour prior to the addition of lactating serum. For each treatment condition, three technical replicate wells were used. One hour prior to the assay, the culture medium was replaced with Agilent Seahorse XF DMEM assay medium supplemented with 1 mM pyruvate, 2 mM glutamine, and 10 mM glucose (pH 7.4). Mitochondrial respiration was measured using the Seahorse XF Cell Mito Stress Test Kit according to the manufacturer’s instructions. Final concentrations of injected compounds were 1 μM oligomycin, 1 μM FCCP (uncoupler), and 0.5 μM rotenone/antimycin A. OCR was measured under basal conditions and following the sequential addition of oligomycin, FCCP, and rotenone/antimycin A. After completion of the Seahorse assay, cells were lysed directly in each well using 0.1% Triton X-100 buffer, and total protein content was quantified using the BCA assay. Basal respiration was calculated by subtracting the average OCR of the final three measurement cycles following rotenone/antimycin A injection (non-mitochondrial respiration) from the average OCR of the first three measurement cycles prior to any inhibitor injection. ATP-coupled respiration was calculated as the difference between basal OCR and OCR following oligomycin injection. Maximal respiration was calculated by subtracting non-mitochondrial OCR from the maximal OCR measured after FCCP injection. All OCR values were normalized to total protein content. Energy maps were generated using Energy Map (Basal rates) with the Seahorse Analytics platform (https://seahorseanalytics.agilent.com). Energy maps were generated using the Seahorse Analytics platform (Agilent) based on basal OCR and ECAR values measured prior to the injection of mitochondrial inhibitors. Specifically, the platform calculates basal rates as the average of the final measurement cycles before oligomycin injection for both OCR and ECAR. All OCR and ECAR values used to generate the energy maps were normalized to protein content (10 ug protein per well).

### Immunostaining

Cultures were fixed with 4% paraformadehyde for 5 minutes at 37 °C and then rinsed with PBS twice for 5 min in 96 well culture plates and blocked for 30 minutes before incubation with anti-H2A.x (EMD Milipore EMD Millipore 05-636) was suspended in blocking buffer (10% horse serum, 2% bovine serum albumin, and 0.5% Triton X-100 in PBS). After wash (10 min PBS plus 0.5% triton X-100, 2×10min PBS), Alexa-flour conjugated secondary antibody (Invitrogen) incubation was performed at a concentration of 1:500 for 1 hour. Samples were then washed again and counterstained with DAPI.

### Mitochondrial staining

A 5 mM stock of MitoSOX Red was freshly prepared in DMEM immediately prior to staining. Live cultures were then rinsed 2x in serum free DMEM before the addition of MitoSOX. Cultures were then incubated for 20 minutes at 37 degrees C in the dark before being rinsed again and imaged with a Axioscope Live imager (Zeiss) under growth conditions. Subsequent images were captured under identical settings.

---

**Supp. Fig. 1: Lactating serum do not affect the growth of breast cancer cells expressing no or low level of PAPP-A**. A) Graph of 3D colony size of indicated breast cancer cells cultured with serum from non-lactating (NL) or lactating mice. B) ELISA of PAPP-A levels in media from the indicated cell lines. ND= not detected.

**Supp. Fig. 2:** List of mitochondrial genes derived from MitoCarta and used for the heat map analysis in figure 2F and figure 3H.

**Supp. Fig. 3:** Table of down-regulated genes driving mitochondrial term hit from RNAseq of MCF7-PAPP-A xenografts.

**Supp. Fig. 4: Lactating serum increases DNA damage**. A) pH2A.X staining of MCF7-PAPP-A cells incubated with non-lactating or lactating serum. B) Quantification of A. Significance was determined by student’s t-test. *p<0.05.

**Supplementary** figure 5: A) List of value obtained for CRF in individual serum samples from non-lactating and lactating women. B) Graph of CRF abundance in lactating versus non-lactating serum.

## ACKNOWLEDGMENTS

**Funding:** This work was supported by the Breast Cancer Research Foundation, The Chemotherapy Foundation and a NCI R21 CA270702-01. We also thank the division of Hematology/Oncology for financial support toward collection of blood samples.

**Authors contributions:** E.C.J has performed all experiments described in this manuscript with the exception of the Seahorse experiments, which were performed by M.C. K.S.A contributed by breeding, collecting and providing serum from mice. S. S helped with the cell culture and analyses. N.T is a clinical research coordinator who wrote to IRB protocol and consented women for this study. J.S is the Chair of Obstetrics and supervised the collection of human samples. C.O provided expertise in PAPP-A biology and supervised the collection of STC1 knockout mice. D.G is the principal investigator who obtained funding and wrote the manuscript.

**Competing interests:** The authors have no conflicting interest to declare.

**Data and methods availability:** All data associated with this study are present in the paper or Supplementary Materials. This study did not generate new unique reagents. The bulk RNA-seq data generated during this study have been deposited in a GEO repository

## Notes

### Competing Interest Statement

The authors have declared no competing interest.

## REFERENCES AND NOTES

1 Collaborative Group on Hormonal Factors in Breast, C. Breast cancer and breastfeeding: collaborative reanalysis of individual data from 47 epidemiological studies in 30 countries, including 50302 women with breast cancer and 96973 women without the disease. Lancet 360, 187–195 (2002). 10.1016/S0140-6736(02)09454-0

2 Victora, C. G., et al. Breastfeeding in the 21st century: epidemiology, mechanisms, and lifelong effect. Lancet 387, 475–490 (2016). 10.1016/S0140-6736(15)01024-7

3 Virassamy, B., et al. Parity and lactation induce T-cell-mediated breast cancer protection. Nature 649, 449–459 (2026). 10.1038/s41586-025-09713-5

4 Little, M., Ye, S. & Fairfax, B. P. Lactation, tissue-resident immunity, and protection against breast cancer. Trends Immunol 47, 3–5 (2026). 10.1016/j.it.2025.12.004

5 Sierra-Roca, J. & Climent, J. Long-Term Breastfeeding: Protective Effects Against Triple-Negative Breast Cancer and the Role of the Breast Microbiota. Pathogens 14 (2025). 10.3390/pathogens14090946

6 Babic, A. et al. Association Between Breastfeeding and Ovarian Cancer Risk. JAMA Oncol 6, e200421 (2020). 10.1001/jamaoncol.2020.0421

7 Eoh, K. J., et al. The preventive effect of breastfeeding against ovarian cancer in BRCA1 and BRCA2 mutation carriers: A systematic review and meta-analysis. Gynecol Oncol 163, 142–147 (2021). 10.1016/j.ygyno.2021.07.028

8 Li, D. P., et al. Breastfeeding and ovarian cancer risk: a systematic review and meta-analysis of 40 epidemiological studies. Asian Pac J Cancer Prev 15, 4829–4837 (2014). 10.7314/apjcp.2014.15.12.4829

9 Augoulea, A., et al. Breastfeeding is associated with lower subclinical atherosclerosis in postmenopausal women. Gynecol Endocrinol 36, 796–799 (2020). 10.1080/09513590.2020.1782374

10 Gunderson, E. P. et al. Lactation Duration and Midlife Atherosclerosis. Obstet Gynecol 126, 381–390 (2015). 10.1097/AOG.0000000000000919

11 Countouris, M. E., et al. Lactation and Maternal Subclinical Atherosclerosis Among Women With and Without a History of Hypertensive Disorders of Pregnancy. J Womens Health (Larchmt) 29, 789–798 (2020). 10.1089/jwh.2019.7863

12 Tschiderer, L., et al. Breastfeeding Is Associated With a Reduced Maternal Cardiovascular Risk: Systematic Review and Meta-Analysis Involving Data From 8 Studies and 1 192 700 Parous Women. J Am Heart Assoc 11, e022746 (2022). 10.1161/JAHA.121.022746

13 Schwarz, E. B., et al. Duration of lactation and risk factors for maternal cardiovascular disease. Obstet Gynecol 113, 974–982 (2009). 10.1097/01.AOG.0000346884.67796.ca

14 Stuebe, A. M. & Rich-Edwards, J. W. The reset hypothesis: lactation and maternal metabolism. Am J Perinatol 26, 81–88 (2009). 10.1055/s-0028-1103034

15 Lawrence, J. B., et al. The insulin-like growth factor (IGF)-dependent IGF binding protein-4 protease secreted by human fibroblasts is pregnancy-associated plasma protein-A. Proc Natl Acad Sci U S A 96, 3149–3153 (1999). 10.1073/pnas.96.6.3149

16 Overgaard, M. T., et al. Pregnancy-associated plasma protein-A2 (PAPP-A2), a novel insulin-like growth factor-binding protein-5 proteinase. J Biol Chem 276, 21849–21853 (2001). 10.1074/jbc.M102191200

17 Boldt, H. B. & Conover, C. A. Pregnancy-associated plasma protein-A (PAPP-A): a local regulator of IGF bioavailability through cleavage of IGFBPs. Growth Horm IGF Res 17, 10–18 (2007). 10.1016/j.ghir.2006.11.003

18 Kalli, K. R., et al. Pregnancy-associated plasma protein-A (PAPP-A) expression and insulin-like growth factor binding protein-4 protease activity in normal and malignant ovarian surface epithelial cells. Int J Cancer 110, 633–640 (2004). 10.1002/ijc.20185

19 Conover, C. A. Key questions and answers about pregnancy-associated plasma protein-A. Trends Endocrinol Metab 23, 242–249 (2012). 10.1016/j.tem.2012.02.008

20 Conover, C. A. & Oxvig, C. The Pregnancy-Associated Plasma Protein-A (PAPP-A) Story. Endocr Rev 44, 1012–1028 (2023). 10.1210/endrev/bnad017

21 Conover, C. A. & Oxvig, C. PAPP-A and cancer. J Mol Endocrinol 61, T1–T10 (2018). 10.1530/JME-17-0236

22 Slocum, E. & Germain, D. Collagen and PAPP-A in the Etiology of Postpartum Breast Cancer. Horm Cancer 10, 137–144 (2019). 10.1007/s12672-019-00368-z

23 Zhang, J., et al. Pregnancy-associated plasma protein-A (PAPPA) promotes breast cancer progression. Bioengineered 13, 291–307 (2022). 10.1080/21655979.2021.2000724

24 Mansfield, A. S., et al. Pregnancy-associated plasma protein-A expression in human breast cancer. Growth Horm IGF Res 24, 264–267 (2014). 10.1016/j.ghir.2014.10.007

25 Poddar, A., et al. The role of pregnancy associated plasma protein-A in triple negative breast cancer: a promising target for achieving clinical benefits. J Biomed Sci 31, 23 (2024). 10.1186/s12929-024-01012-x

26 Prithviraj, P., et al. Aberrant pregnancy-associated plasma protein-A expression in breast cancers prognosticates clinical outcomes. Sci Rep 10, 13779 (2020). 10.1038/s41598-020-70774-9

27 Hjortebjerg, R., Hogdall, C., Hansen, K. H., Hogdall, E. & Frystyk, J. The IGF-PAPP-A-Stanniocalcin Axis in Serum and Ascites Associates with Prognosis in Patients with Ovarian Cancer. Int J Mol Sci 25 (2024). 10.3390/ijms25042014

28 Thomsen, J., et al. PAPP-A proteolytic activity enhances IGF bioactivity in ascites from women with ovarian carcinoma. Oncotarget 6, 32266–32278 (2015). 10.18632/oncotarget.5010

29 Becker, M. A., Haluska, P., Jr., Bale, L. K., Oxvig, C. & Conover, C. A. A novel neutralizing antibody targeting pregnancy-associated plasma protein-a inhibits ovarian cancer growth and ascites accumulation in patient mouse tumorgrafts. Mol Cancer Ther 14, 973–981 (2015). 10.1158/1535-7163.MCT-14-0880

30 Boldt, H. B. & Conover, C. A. Overexpression of pregnancy-associated plasma protein-A in ovarian cancer cells promotes tumor growth in vivo. Endocrinology 152, 1470–1478 (2011). 10.1210/en.2010-1095

31 Steffensen, L. B., Conover, C. A. & Oxvig, C. PAPP-A and the IGF system in atherosclerosis: what’s up, what’s down? Am J Physiol Heart Circ Physiol 317, H1039–H1049 (2019). 10.1152/ajpheart.00395.2019

32 Conover, C. A. Pregnancy-associated plasma protein-A (PAPP-A) and cardiovascular disease. Growth Horm IGF Res 79, 101625 (2024). 10.1016/j.ghir.2024.101625

33 Consuegra-Sanchez, L., Fredericks, S. & Kaski, J. C. Pregnancy-associated plasma protein-A (PAPP-A) and cardiovascular risk. Atherosclerosis 203, 346–352 (2009). 10.1016/j.atherosclerosis.2008.07.042

34 Yu, X. H., et al. Pregnancy-associated plasma protein-A in atherosclerosis: Molecular marker, mechanistic insight, and therapeutic target. Atherosclerosis 278, 250–258 (2018). 10.1016/j.atherosclerosis.2018.10.004

35 Takabatake, Y., et al. Lactation opposes pappalysin-1-driven pregnancy-associated breast cancer. EMBO Mol Med 8, 388–406 (2016). 10.15252/emmm.201606273

36 Heitzeneder, S., et al. Pregnancy-Associated Plasma Protein-A (PAPP-A) in Ewing Sarcoma: Role in Tumor Growth and Immune Evasion. J Natl Cancer Inst 111, 970–982 (2019). 10.1093/jnci/djy209

37 Jenkins, E. C., Brown, S. O. & Germain, D. The Multi-Faced Role of PAPP-A in Post-Partum Breast Cancer: IGF-Signaling is Only the Beginning. J Mammary Gland Biol Neoplasia 25, 181–189 (2020). 10.1007/s10911-020-09456-1

38 Kirschner, A., et al. Pappalysin-1 T cell receptor transgenic allo-restricted T cells kill Ewing sarcoma in vitro and in vivo. Oncoimmunology 6, e1273301 (2017). 10.1080/2162402X.2016.1273301

39 Conover, C. A., Bale, L. K. & Nair, K. S. Comparative gene expression and phenotype analyses of skeletal muscle from aged wild-type and PAPP-A-deficient mice. Exp Gerontol 80, 36–42 (2016). 10.1016/j.exger.2016.04.005

40 Alassaf, M. & Halloran, M. C. Pregnancy-associated plasma protein-aa regulates endoplasmic reticulum-mitochondria associations. Elife 10 (2021). 10.7554/eLife.59687

41 Bhardwaj, G., et al. Transcriptomic Regulation of Muscle Mitochondria and Calcium Signaling by Insulin/IGF-1 Receptors Depends on FoxO Transcription Factors. Front Physiol 12, 779121 (2021). 10.3389/fphys.2021.779121

42 Sadaba, M. C., Martin-Estal, I., Puche, J. E. & Castilla-Cortazar, I. Insulin-like growth factor 1 (IGF-1) therapy: Mitochondrial dysfunction and diseases. Biochim Biophys Acta 1862, 1267–1278 (2016). 10.1016/j.bbadis.2016.03.010

43 Biswas, S., Ghosh, S. & Maitra, S. Role of insulin-like growth factor 1 (IGF1) in the regulation of mitochondrial bioenergetics in zebrafish oocytes: lessons from in vivo and in vitro investigations. Front Cell Dev Biol 11, 1202693 (2023). 10.3389/fcell.2023.1202693

44 Guan, X., Yan, Q., Wang, D., Du, G. & Zhou, J. IGF-1 Signaling Regulates Mitochondrial Remodeling during Myogenic Differentiation. Nutrients 14 (2022). 10.3390/nu14061249

45 Katsenelson, M., et al. IGF-1 receptor regulates upward firing rate homeostasis via the mitochondrial calcium uniporter. Proc Natl Acad Sci U S A 119, e2121040119 (2022). 10.1073/pnas.2121040119

46 Glover, H. L., Schreiner, A., Dewson, G. & Tait, S. W. G. Mitochondria and cell death. Nat Cell Biol 26, 1434–1446 (2024). 10.1038/s41556-024-01429-4

47 Blume, A., et al. Prolactin activates mitogen-activated protein kinase signaling and corticotropin releasing hormone transcription in rat hypothalamic neurons. Endocrinology 150, 1841–1849 (2009). 10.1210/en.2008-1023

48 Brunton, P. J., Russell, J. A. & Douglas, A. J. Adaptive responses of the maternal hypothalamic-pituitary-adrenal axis during pregnancy and lactation. J Neuroendocrinol 20, 764–776 (2008). 10.1111/j.1365-2826.2008.01735.x

49 Gustafson, P. E., et al. The role of prolactin in the suppression of the response to restraint stress in the lactating mouse. J Neuroendocrinol 36, e13330 (2024). 10.1111/jne.13330

50 Gustafson, P., Kokay, I., Sapsford, T., Bunn, S. & Grattan, D. Prolactin regulation of the HPA axis is not mediated by a direct action upon CRH neurons: evidence from the rat and mouse. Brain Struct Funct 222, 3191–3204 (2017). 10.1007/s00429-017-1395-1

51 Paul, N., et al. Human placenta releases extracellular vesicles carrying corticotrophin releasing hormone mRNA into the maternal blood. Placenta 146, 71–78 (2024). 10.1016/j.placenta.2024.01.004

52 Lightman, S. L., et al. Peripartum plasticity within the hypothalamo-pituitary-adrenal axis. Prog Brain Res 133, 111–129 (2001). 10.1016/s0079-6123(01)33009-1

53 Bosch, O. J., Musch, W., Bredewold, R., Slattery, D. A. & Neumann, I. D. Prenatal stress increases HPA axis activity and impairs maternal care in lactating female offspring: implications for postpartum mood disorder. Psychoneuroendocrinology 32, 267–278 (2007). 10.1016/j.psyneuen.2006.12.012

54 Battaglia, C. R., Cursano, S., Calzia, E., Catanese, A. & Boeckers, T. M. Corticotropin-releasing hormone (CRH) alters mitochondrial morphology and function by activating the NF-kB-DRP1 axis in hippocampal neurons. Cell Death Dis 11, 1004 (2020). 10.1038/s41419-020-03204-3

55 Isola, R., Solinas, P., Loy, F., Mariotti, S. & Riva, A. 3-D structure of mitochondrial cristae in rat adrenal cortex varies after acute stimulation with ACTH and CRH. Mitochondrion 10, 472–478 (2010). 10.1016/j.mito.2010.05.007

56 Zhang, Y., et al. Stress triggers gut dysbiosis via CRH-CRHR1-mitochondria pathway. NPJ Biofilms Microbiomes 10, 93 (2024). 10.1038/s41522-024-00571-z

57 Bornstein, S. R., Ehrhart-Bornstein, M., Guse-Behling, H. & Scherbaum, W. A. Structure and dynamics of adrenal mitochondria following stimulation with corticotropin releasing hormone. Anat Rec 234, 255–262 (1992). 10.1002/ar.1092340212

58 Graziani, G., et al. Evidence that corticotropin-releasing hormone inhibits cell growth of human breast cancer cells via the activation of CRH-R1 receptor subtype. Mol Cell Endocrinol 264, 44–49 (2007). 10.1016/j.mce.2006.10.006

59 Stuhr, L. E., Wei, E. T. & Reed, R. K. Corticotropin-releasing factor reduces tumor volume, halts further growth, and enhances the effect of chemotherapy in 4T1 mammary carcinoma in mice. Tumour Biol 35, 1365–1370 (2014). 10.1007/s13277-013-1186-0

60 Graziani, G., et al. CRH inhibits cell growth of human endometrial adenocarcinoma cells via CRH-receptor 1-mediated activation of cAMP-PKA pathway. Endocrinology 143, 807–813 (2002). 10.1210/endo.143.3.8694

61 Tjuvajev, J., et al. Anti-neoplastic properties of human corticotropin releasing factor: involvement of the nitric oxide pathway. In Vivo 12, 1–10 (1998).

62 Sato, N., et al. Expression of Corticotropin-Releasing Hormone and Its Receptors May Be Associated With Survival Rate in Pancreatic Cancer. Gastro Hep Adv 2, 147–155 (2023). 10.1016/j.gastha.2022.09.003

63 Jin, L., et al. Corticotropin-releasing hormone receptors mediate apoptosis via cytosolic calcium-dependent phospholipase A(2) and migration in prostate cancer cell RM-1. J Mol Endocrinol 52, 255–267 (2014). 10.1530/JME-13-0270

64 Moliterno, J. A., Henry, E. & Pannullo, S. C. Corticorelin acetate injections for the treatment of peritumoral brain edema. Expert Opin Investig Drugs 18, 1413–1419 (2009). 10.1517/13543780903190689

65 Bischof, P. Purification and characterization of pregnancy associated plasma protein A (PAPP-A). Arch Gynecol 227, 315–326 (1979). 10.1007/BF02109920

66 Costanzo, V., Bardelli, A., Siena, S. & Abrignani, S. Exploring the links between cancer and placenta development. Open Biol 8 (2018). 10.1098/rsob.180081

67 Rekowska, A. K., et al. Biomolecules Involved in Both Metastasis and Placenta Accreta Spectrum-Does the Common Pathophysiological Pathway Exist? Cancers (Basel) 15 (2023). 10.3390/cancers15092618

68 Hossain, S. M., Lynch-Sutherland, C. F., Chatterjee, A., Macaulay, E. C. & Eccles, M. R. Can Immune Suppression and Epigenome Regulation in Placenta Offer Novel Insights into Cancer Immune Evasion and Immunotherapy Resistance? Epigenomes 5 (2021). 10.3390/epigenomes5030016

69 Lala, P. K., Nandi, P., Hadi, A. & Halari, C. A crossroad between placental and tumor biology: What have we learnt? Placenta 116, 12–30 (2021). 10.1016/j.placenta.2021.03.003

70 Dewerchin, M. & Carmeliet, P. Placental growth factor in cancer. Expert Opin Ther Targets 18, 1339–1354 (2014). 10.1517/14728222.2014.948420

71 Mao, X., et al. Association of reproductive risk factors and breast cancer molecular subtypes: a systematic review and meta-analysis. BMC Cancer 23, 644 (2023). 10.1186/s12885-023-11049-0

72 Ambrosone, C. B. & Higgins, M. J. Relationships between Breast Feeding and Breast Cancer Subtypes: Lessons Learned from Studies in Humans and in Mice. Cancer Res 80, 4871–4877 (2020). 10.1158/0008-5472.CAN-20-0077

73 Elias, K. M., Berkowitz, R. S. & Horowitz, N. S. Ultra High-risk Gestational Trophoblastic Neoplasia. Hematol Oncol Clin North Am 38, 1259–1264 (2024). 10.1016/j.hoc.2024.08.015

74 Baas, I. O., et al. Immunotherapy for Gestational Trophoblastic Neoplasia: A New Paradigm. Gynecol Obstet Invest 89, 230–238 (2024). 10.1159/000533972

75 Conover, C. A., Bale, L. K. & Oxvig, C. Targeted Inhibition of Pregnancy-Associated Plasma Protein-A Activity Reduces Atherosclerotic Plaque Burden in Mice. J Cardiovasc Transl Res 9, 77–79 (2016). 10.1007/s12265-015-9666-9

76 Bale, L. K., Chakraborty, S. & Conover, C. A. Inducible reduction in pregnancy-associated plasma protein-A gene expression inhibits established atherosclerotic plaque progression in mice. Endocrinology 155, 1184–1187 (2014). 10.1210/en.2013-2110

77 Recht, L., Mechtler, L. L., Wong, E. T., O’Connor, P. C. & Rodda, B. E. Steroid-sparing effect of corticorelin acetate in peritumoral cerebral edema is associated with improvement in steroid-induced myopathy. J Clin Oncol 31, 1182–1187 (2013). 10.1200/JCO.2012.43.9455

78 Ibanez, L., Potau, N., Marcos, M. V. & de Zegher, F. Corticotropin-releasing hormone: a potent androgen secretagogue in girls with hyperandrogenism after precocious pubarche. J Clin Endocrinol Metab 84, 4602–4606 (1999). 10.1210/jcem.84.12.6239

79 Wu, Y., et al. Enhanced expression of vascular cell adhesion molecule-1 by corticotrophin-releasing hormone contributes to progression of atherosclerosis in LDL receptor-deficient mice. Atherosclerosis 203, 360–370 (2009). 10.1016/j.atherosclerosis.2008.05.059

80 Henein, M. Y., Vancheri, S., Longo, G. & Vancheri, F. The Impact of Mental Stress on Cardiovascular Health-Part II. J Clin Med 11 (2022). 10.3390/jcm11154405

81 Slocum, E., Craig, A., Villanueva, A. & Germain, D. Parity predisposes breasts to the oncogenic action of PAPP-A and activation of the collagen receptor DDR2. Breast Cancer Res 21, 56 (2019). 10.1186/s13058-019-1142-z

82 Peterson, E. A., et al. Amphiregulin Is a Critical Downstream Effector of Estrogen Signaling in ERalpha-Positive Breast Cancer. Cancer Res 75, 4830–4838 (2015). 10.1158/0008-5472.CAN-15-0709

83 Torre, D., Lachmann, A. & Ma’ayan, A. BioJupies: Automated Generation of Interactive Notebooks for RNA-Seq Data Analysis in the Cloud. Cell Syst 7, 556–561 e553 (2018). 10.1016/j.cels.2018.10.007

